# Navigating Correlation, Causation, and Interpretation in Resting-State fMRI

**DOI:** 10.1101/2025.09.16.676611

**Authors:** Gang Chen, Zhengchen Cai, Konrad P. Kording, Thomas T. Liu, Joshua Faskowitz, Peter A. Bandettini, Bharat Biswal, Paul A. Taylor

## Abstract

Resting-state fMRI has generated influential insights into large-scale brain organization and contributed to clinically relevant applications, largely through correlation-based measures of cross-regional association in BOLD responses. At the same time, interpreting these statistical associations as reflecting underlying neural interactions requires careful consideration of fundamental methodological constraints.

Here, we distinguish two fundamental but often conflated limitations. The first is *measurement distortion*: the fMRI signal is an indirect and heterogeneous measurement of neural activity, arising from neurovascular coupling, physiology, and measurement-related processes, which can introduce systematic and incompletely characterized biases into estimated correlations. The second is *causal non-identifiability*: even if correlations perfectly reflected neural synchrony, the resulting correlation structure would not uniquely determine the underlying neural interactions.

Using causal reasoning, simulations, and analytic arguments, we distinguish these limitations and examine their consequences for interpretation. Although both apply broadly to fMRI, their implications are particularly important in resting-state analyses, where correlation structure is the primary object of inference in the absence of experimental perturbation. We show that measurement-related biases can distort estimated correlations (e.g., attenuating or inflating associations and affecting group comparisons) and can produce reproducible patterns that do not necessarily reflect underlying neural relationships, highlighting that statistical reliability does not guarantee biological validity. We further show that graph-theoretic, geometric, and other higher-order representations derived from these correlations do not, by themselves, justify mechanistic interpretation. We argue not against the utility of resting-state fMRI, but for greater precision in interpretation. Our conclusions concern the interpretation of correlation-based analyses rather than their methodological utility. Distinguishing measurement distortion from causal non-identifiability clarifies the inferential boundaries separating descriptive association, predictive utility, and mechanistic interpretation.

## 1 Introduction

Resting-state functional magnetic resonance imaging (fMRI) has become a central tool for characterizing large-scale brain organization. Correlation-based measures between fMRI time series, commonly referred to as *functional connectivity* (Friston, 2011), have revealed reproducible network patterns such as the default mode network and informed studies of neurological and psychiatric disorders (e.g., Biswal et al., 1995; Greicius et al., 2003; Fox et al., 2009; Margulies et al., 2016; Sporns, 2018; Mitra et al., 2023). These approaches have also contributed to translational applications, including neuromodulation targeting and biomarker development (e.g., Fox, 2010; Castellanos et al., 2013; Drysdale et al., 2016; Hopman et al., 2021). Importantly, correlation-based associations reflect statistical associations in blood-oxygenation-level-dependent (BOLD) signals rather than direct measures of neural interaction. Nonetheless, correlation-based fMRI analyses have demonstrated robust descriptive and predictive value, yielding clinically meaningful insights even while operating at a level distinct from underlying neuronal processes.

### 1.1 Target of inference in resting-state fMRI

In resting-state fMRI, BOLD correlations are the primary quantities of interest and are typically summarized as correlation matrices or network representations (Fig. 1). A central but often implicit question concerns the target of inference for these matrices. While correlations are routinely treated as proxies for cross-regional neural relationships, the intended underlying quantity—whether synchronous neural activity, direct or indirect interaction, shared inputs, or broader organizational structure—is rarely specified. Because the fMRI signal is an indirect and heterogeneous measurement, and because correlation does not uniquely map to causal relationships, the link between observed associations and their neural interpretation remains fundamentally ambiguous. Clarifying this target is essential for determining what correlation-based analyses can and cannot reveal.

**Figure 1:**
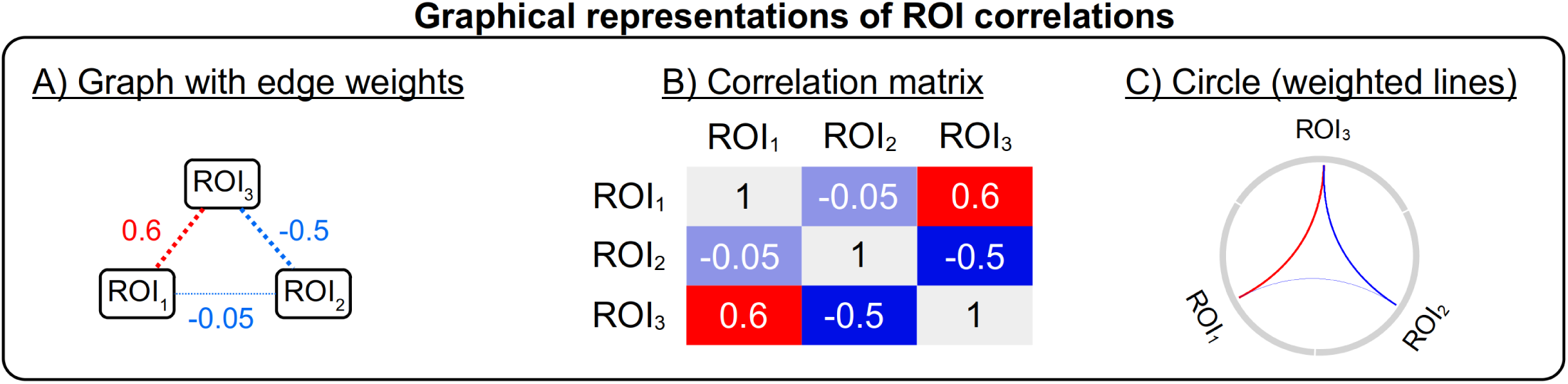
Correlation structure among three regions of interest (ROIs). (A) Pairwise associations as a triangle: dotted lines denote correlations. (B) Corresponding 3 × 3 correlation matrix. (C) Circos plot visualization: arc thickness reflects correlation magnitude and color (red/blue) indicates sign. These representations show that resting-state analyses operate at the level of pairwise association, without specifying directionality.

When estimated correlations are further transformed into higher-order representations of correlation structure—including graph-theoretic metrics (e.g., node degree, hubness, rich club, community structure) (Sporns, 2018), gradient-based embeddings (Margulies et al., 2016), and geometric or topological summaries (Pang et al., 2023)—the same interpretational ambiguities persist. Because these macroscale metrics are deterministic functions of the underlying correlations, they inherit the same assumptions and potential biases. Thus, even if correlations accurately reflected neural-level associations, interpreting such metrics as evidence of direct neural communication or information flow requires causal assumptions far beyond what the statistical data alone can support.

### 1.2 General constraints inherent to fMRI through the lens of causal inference

To formalize these relationships, we use directed acyclic graphs (DAGs) (Pearl, 2009), which provide a transparent framework for representing assumptions about how neural activity, physiological processes, and measurement factors jointly give rise to observed fMRI signals (Fig. 2). In resting-state fMRI, no variable is experimentally manipulated, and observed correlations reflect a mixture of direct neural interactions, shared inputs, and non-neural influences. This causal representation makes explicit that observed data arise from a mixture of neural, physiological, and measurement-related influences, rather than from neural interactions alone, highlighting that observed associations do not uniquely reflect neural interactions.

**Figure 2:**
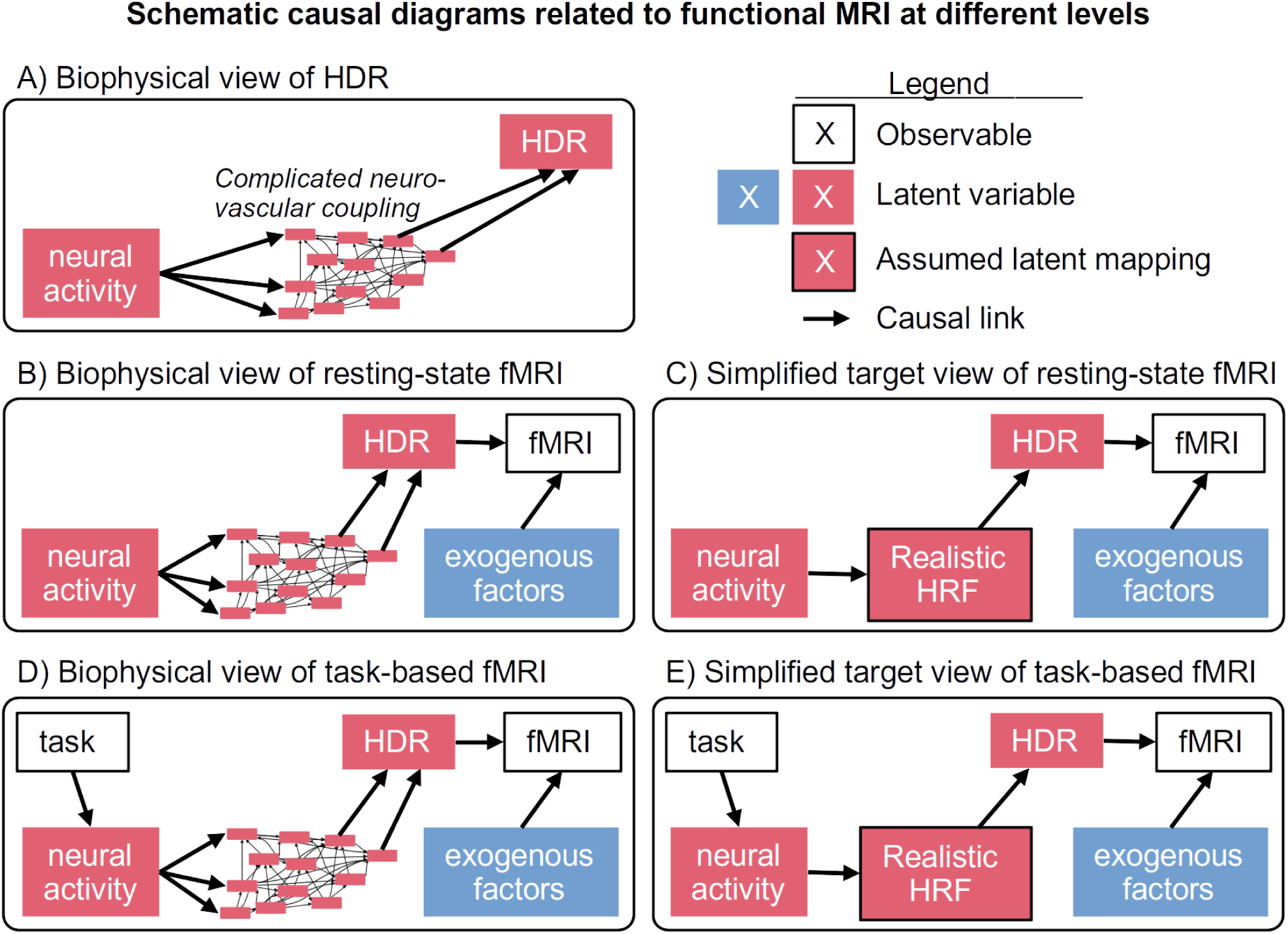
Directed acyclic graph (DAG) representations of relationships among neural activity, the hemodynamic response (HDR), and the fMRI (BOLD) signal. Arrows denote assumed causal relations; colored boxes indicate unobserved processes. (A) **Biophysical mediation.** Neural activity influences the HDR through neurovascular coupling. (B) **Resting-state.** Observed fMRI signals reflect neural activity together with physiological and measurement-related influences. (C) HRF abstraction of neurovascular coupling. (D–E) **Task-based.** Experimental manipulation introduces an intervention that anchors BOLD-level inference.

We distinguish two conceptually distinct but often conflated limitations inherent to fMRI:

- **Measurement Distortion:** A problem of estimation accuracy. The fMRI signal is an indirect and heterogeneous transformation of neural activity, shaped by variable neurovascular coupling, physiological processes, and measurement artifacts (Logothetis, 2008). These unmodeled influences can introduce systematic biases, causing estimated statistical associations to substantially under- or over-represent the underlying neural synchrony.
- **Causal Non-identifiability:** A problem of inference. Even if measurement distortion were eliminated, the resulting correlation structure does not uniquely determine the underlying neural interactions. Distinct causal architectures can generate identical correlation matrices.

These two limitations operate at different levels: measurement distortion affects how accurately correlations are estimated, whereas causal non-identifiability limits what can be inferred from them even if measured without error.

This DAG perspective has an important implication: different underlying causal structures can give rise to similar patterns of statistical association. Consequently, correlations—and quantities derived from them, including graph-theoretic metrics—do not uniquely determine neural interactions, distinguish direct from indirect effects, or establish directionality. We examine these limitations formally in Sections 2 and 3.

### 1.3 Task vs. rest: why unanchored correlations are especially vulnerable

While these limitations apply broadly across fMRI, their consequences differ between resting-state and task-based paradigms. Three related distinctions clarify this difference. First, the target of inference differs. In task-based fMRI, experimental manipulation allows inference to focus on changes in BOLD response magnitude, often expressed in physically interpretable units such as percent signal change. Within a specified modeling framework, these quantities can be interpreted as reflecting the effect of the task condition on the measured signal at the BOLD level. In contrast, resting-state fMRI is typically characterized by correlations between regions, which are dimensionless measures of statistical association. Unlike task-evoked responses, these quantities do not correspond to a directly manipulated variable. As a result, while they are well defined as summaries of BOLD-level association, their interpretation as reflecting underlying neural interactions requires additional assumptions or constraints and is not uniquely supported by the data alone.

Second, the paradigms differ in how inference is anchored. Task-based fMRI incorporates experimentally imposed manipulations that introduce externally controlled variation (Fig. 2D,E). This variation serves as an inferential anchor, allowing time-series models to relate changes in fMRI signal to known inputs. In typical analyses, task timing is encoded through regressors constructed via convolution with a hemodynamic response function (HRF). Within this framework, exogenous noise (motion, breathing, cardiac pulsation, scanner distortions, etc.) can be partially separated from task-related signals, and modeling neurovascular coupling—especially with flexible HRFs tailored to regions, tasks, or individuals (Prince et al., 2022; Chen et al., 2023)—can improve the accuracy of estimated task-evoked BOLD responses. This separation is further supported by experimental design features such as randomization of trial conditions and temporal jittering, which introduce variability that is independent of ongoing physiological and measurement-related processes, thereby reducing systematic coupling between task effects and non-neural variability. Although this separation is not complete, it provides an important inferential advantage.

In contrast, resting-state fMRI is purely observational, lacking experimentally imposed variation to anchor causal interpretation. As a result, and typically without explicit modeling of neurovascular coupling, resting-state analyses lack a comparable mechanism for separating exogenous noise from the mediated effects of heterogeneous neurovascular processes (Fig. 2B,C). Because the correlation structure itself is the primary object of inference, these influences are absorbed directly into cross-region correlation estimates rather than explicitly modeled. Consequently, correlations may be over- or under-estimated, and group comparisons may be attenuated, inflated, or even sign-reversed (Section 2). Importantly, this distinction applies already at the level of the measured signal. Even when interpretation is restricted to BOLD dynamics, resting-state correlations alone do not identify causal structure and therefore require additional assumptions or external constraints for causal interpretation because measurement distortion and non-identifiability prevent unique attribution of observed associations.

Third, the paradigms differ in how the mapping from neural activity to the BOLD signal is modeled. This structural difference does not fully eliminate measurement distortion in task-based fMRI, but it provides an external reference that constrains interpretation, whereas no such constraint exists in resting-state analyses. This distinction does not remove broader challenges in modeling the mapping between neural activity and fMRI signal (Epp et al., 2025; Goltermann et al., 2026; Proulx and Farivar, 2026). In task-based fMRI, the HRF is explicitly incorporated into the modeling framework, allowing the mapping from neural activity to fMRI signal to be estimated, albeit imperfectly. In resting-state analyses, by contrast, this mapping is typically not modeled explicitly; its variability is absorbed into observed correlations, contributing to measurement distortion. As a result, resting-state analyses must contend simultaneously with unpartitioned noise and the absence of an external anchor for interpretation. Recent work comparing resting-state and multi-task datasets further suggests that sampling neural activity across diverse task contexts can improve estimation of functional organization (Nettekoven et al., 2026).

These distinctions are summarized in Table 1.

**Table 1:** Structural differences between task-based and resting-state fMRI that affect inferential properties.

| Aspect | Task-based fMRI | Resting-state fMRI |
| --- | --- | --- |
| *Experimental design and measurement structure* |  |  |
| Experimental manipulation | Yes—externally imposed variation provides an inferential anchor | No—purely observational |
| Design structure | Randomization and temporal jittering help separate signal from noise | No decoupling of signal and noise from experimental design |
| Noise disentanglement | Partial separation of task-related signal from exogenous noise via modeling | Noise contributions are absorbed directly into correlation estimates |
| *Target of inference and interpretation* |  |  |
| Quantity of inference | BOLD response magnitude (or profile) | Cross-region correlation |
| Nature of quantity | Physically interpretable (e.g., percent signal change) | Dimensionless statistical association |
| Implications for inference | Measurement distortion partially constrained by design and modeling | Measurement distortion directly reflected in estimated correlations |
| Causal interpretability | Causal inference of task effects on BOLD can be supported under model assumptions | Correlation structure alone does not identify causal structure; causal interpretation requires additional assumptions or external constraints |

### 1.4 Operationalizing the levels of inference

Our aim is not to diminish the utility of resting-state fMRI, nor to suggest that correlation-based analyses are uninformative. Rather, interpretation must align with the level of inference supported by the data. We distinguish three levels:

(1) **Descriptive Association:** Statements that regions exhibit correlated BOLD responses are supported as summaries of measured signals, though estimates may be biased by measurement distortion (Section 2). Such distortions can propagate to group-level analyses, potentially producing spurious differences, masking true effects, or even reversing the direction of genuine differences.
(2) **Predictive/Biomarker Utility:** Reproducible correlation patterns can be effectively used for classification, machine learning, or clinical intervention targeting, even when influenced by stable non-neural factors. Even for predictive or biomarker applications, however, measurement distortion and causal non-identifiability can affect the stability, interpretability, and generalizability of derived features.
(3) **Mechanistic/Causal Interaction:** Claims about directionality, direct communication, or information flow are not identifiable from resting-state correlations alone and require additional assumptions or experimental evidence (Section 3).

### 1.5 Overview

Because correlation is the primary target of inference in resting-state fMRI, evaluating the accuracy of its estimation is critical. Statistical estimation involves both precision (uncertainty) and accuracy (bias); however, in neuroimaging, the high reproducibility of correlation patterns is often conflated with biological validity. Reproducibility primarily reflects statistical stability, meaning that even highly reproducible correlations may systematically misrepresent underlying neural relationships.

This raises a central question: to what extent do estimated correlations provide an accurate proxy for neural-level associations, and how do inherent biases affect downstream inferences, such as group comparisons and graph-theoretic metrics of network topology? Using causal reasoning, simulations, and analytic arguments, we address three questions:

(1) How accurately do BOLD correlations capture cross-regional neural relationships in the presence of unpar-titioned noise?
(2) What are the consequences of inaccuracies in estimated correlations on group-level inferences?
(3) Under what assumptions can correlations support mechanistic interpretation?

Distinguishing measurement distortion from causal non-identifiability provides a unified framework for evaluating correlation-based analyses and clarifies the boundary between descriptive association and mechanistic inference. These considerations extend to many downstream procedures that operate on correlation structure, including graph-theoretic analyses, gradient-based embeddings, feature-selection approaches, and other methods that transform or sparsify correlation matrices.

All simulation code is available at: https://github.com/afni/apaper_DAG2.

## 2 Measurement distortion in resting-state fMRI

Measurement distortion concerns how accurately correlations estimated from fMRI signals reflect underlying neural relationships. Because the fMRI signal is an indirect and heterogeneous transformation of neural activity (Fig. 2A–C), estimated correlations may deviate systematically from neural-level associations. This section focuses on how properties of the measurement process, independent of sampling variability, affect correlation estimates.

Measurement distortion directly challenges three common heuristic assumptions used when interpreting correlation matrices (Fig. 1):

(1) that near-zero correlation implies an absence of interaction;
(2) that the sign of correlation reflects excitatory (+) or inhibitory (−) influence; and
(3) that correlation magnitude reflects underlying neural interaction strength.

To illustrate how these assumptions can fail at the measurement level, we consider the combined effects of two inseparable sources of distortion: exogenous noise and unmodeled neurovascular coupling. The simulations presented below demonstrate general mathematical properties of correlation under plausible measurement structures, highlighting how distortion structurally arises even under simplified conditions.

### 2.1 Noise-induced bias in correlation estimates: individual level

Resting-state fMRI relies on correlations between fMRI signals to characterize cross-regional synchrony. The simulations below examine how properties of the measurement process influence these estimated correlations. For clarity, key symbols are defined in Table 3, while the full analytical relationships between neural and BOLD correlations are provided in Appendix A.

**Case 1: fMRI correlation in the presence of simple noise.** We first consider a simplified scenario in which noise is independent of neural activity and uncorrelated across regions. Here, noise *ɛ_i_* represents exogenous influences such as motion, physiological fluctuations, and scanner variability. In the corresponding causal structure (Fig. 3), the observed correlation *r*_fMRI_ serves as a proxy for the latent neural correlation *r*_neur_ . The neural component HDR.neural*_i_* represents the neurally driven hemodynamic response in region *i* under an ideal mapping from neural activity, such that the neural correlation *r*_neur_ would be recovered without distortion. Simulation results (Fig. 4) show that as noise increases relative to the neural signal, *r*_fMRI_ is systematically attenuated toward zero. For example, when noise and signal have comparable magnitude, correlations may be reduced by roughly 50%; when noise dominates, correlations approach zero, obscuring genuine neural relationships. This attenuation effect is well known in classical statistics (Spearman, 1904) and is consistent with empirical observations that individual-level resting-state correlations are typically modest in magnitude (e.g., 0.0–0.3 for cortex–cortex pairs and 0.0–0.2 for subcortical–cortical pairs) (Luckett et al., 2023).

**Figure 3:**
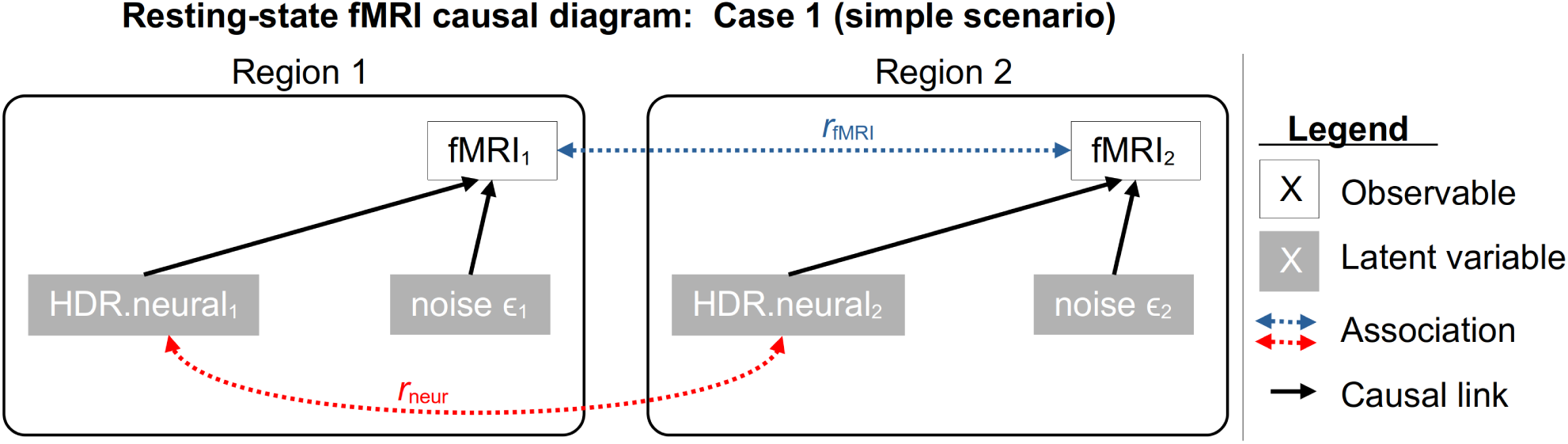
Causal diagram illustrating a two-region scenario in resting-state fMRI. The observed correlation *r*_fMRI_ is sys-tematically distorted relative to the latent neural correlation *r*_neur_ due to noise contributions.

**Figure 4:**
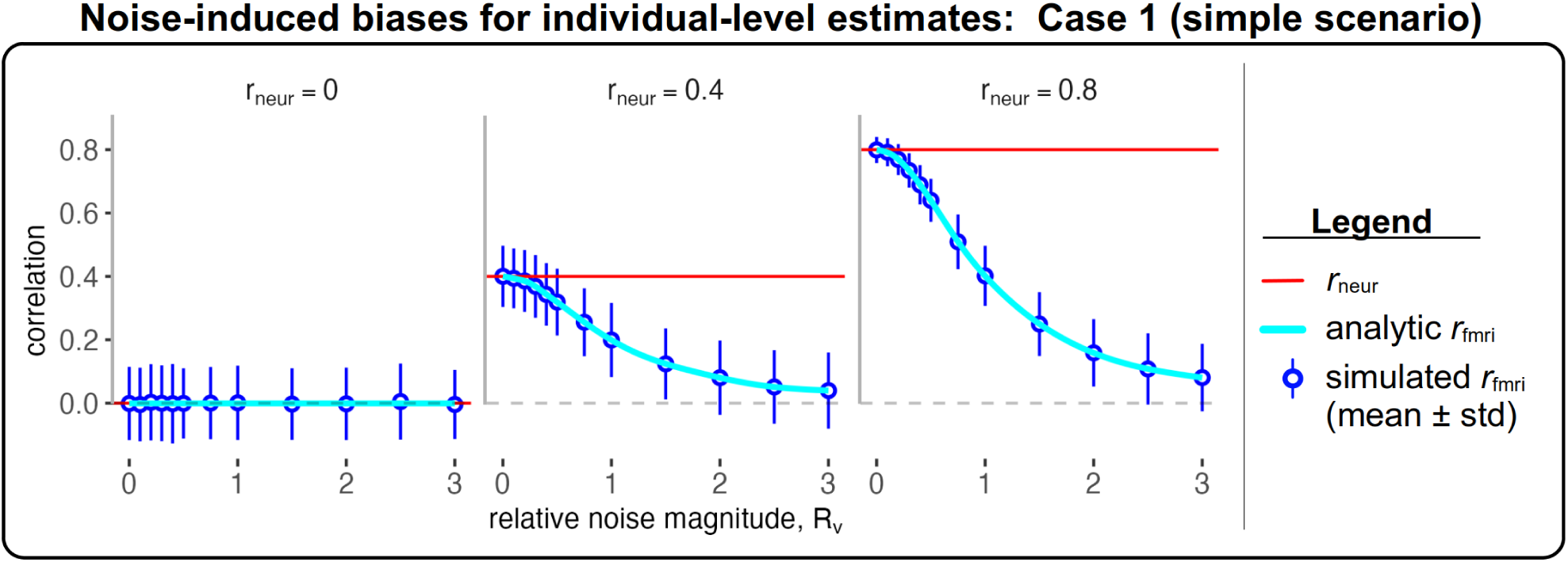
Simulation results illustrating attenuation of *r*_fMRI_ relative to *r*_neur_ under increasing noise levels. Analytical framework for the simulations is described in Appendix A.

These results illustrate a baseline effect of independent noise. However, real-world fMRI signals are influenced by more complex noise structures, including shared noise sources, region-specific noise levels, and variability in the latency, amplitude, and shape of both neural and hemodynamic responses. As these realistic aspects are incorporated, their heterogeneity distorts correlation estimates in more intricate ways.

**Case 2: fMRI correlation in the presence of structured noise.** We next consider a more realistic setting in which noise is correlated with neural signals and across regions. Deviations from an ideal neural-to-BOLD mapping, including nonlinearities and regional neurovascular differences, are absorbed into a residual term *ɛ_i_*. Here, the term “noise” is used broadly to denote variability not explicitly modeled at the neural level. This includes two sources of variability: (i) exogenous influences, such as motion, physiological fluctuations, and scanner-related artifacts, which are external to the neural signal; and (ii) endogenous variability arising from neurovascular coupling, which reflects nonlinear and spatially heterogeneous transformations from neural activity to the observed BOLD response. This endogenous variability implies that the mapping between neural activity and BOLD responses is not uniform across regions, and may spatially vary with vascular properties and metabolic demands (Ekstrom, 2020).

This formulation introduces two additional dependencies: neural–noise mixing (*ρ*_mix_) and noise–noise correla-tion (*ρ_ɛ_*) (Fig. 5). These dependencies naturally arise from heterogeneous neurovascular coupling and acquisition-related spatial structure (Kellman and McVeigh, 2005). For simplicity, *ρ*_mix_ is assumed to be fixed, effectively treating the unmodeled hemodynamic mapping as homogeneous across regions. While biologically unrealistic, it provides a useful mathematical baseline for understanding how noise–signal coupling influences observed correlations.

**Figure 5:**
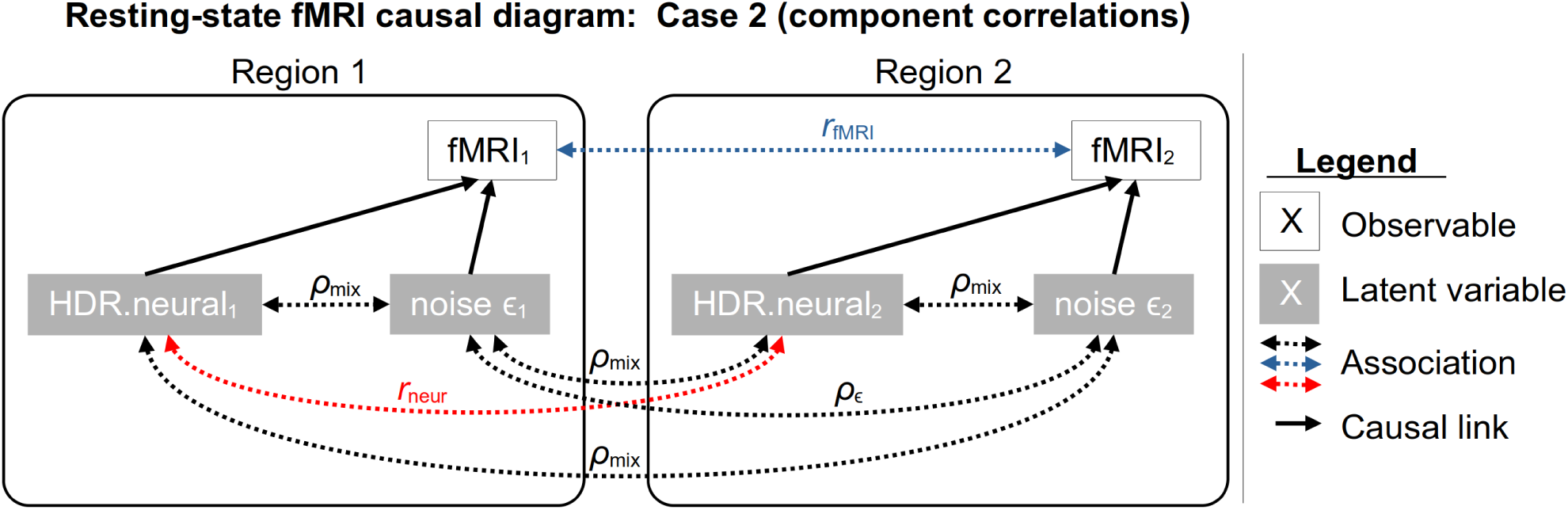
Causal diagram incorporating neural–noise mixing and shared noise.

Simulation results (Fig. 6) show that these dependencies produce more complex bias patterns. Correlations may be attenuated or inflated, depending on the interaction between signal and noise. For example, nonzero *ρ_ɛ_* can induce positive correlations even when *r*_neur_ = 0, while neural–noise coupling can either offset attenuation or introduce additional bias. Furthermore, when *ρ*_mix_ = 0 (Fig. 6A), *r*_fMRI_ estimates are biased toward *ρ_ɛ_*, as previously noted by (Liu, 2013). However, when *ρ*_mix_ ≠ 0 (Fig. 6B), the direction of bias varies with the noise magnitude: when neural activity and noise are correlated, cross-regional noise synchrony can partially offset underestimation or induce artifactual correlations even when *r*_neur_ = 0.

**Figure 6:**
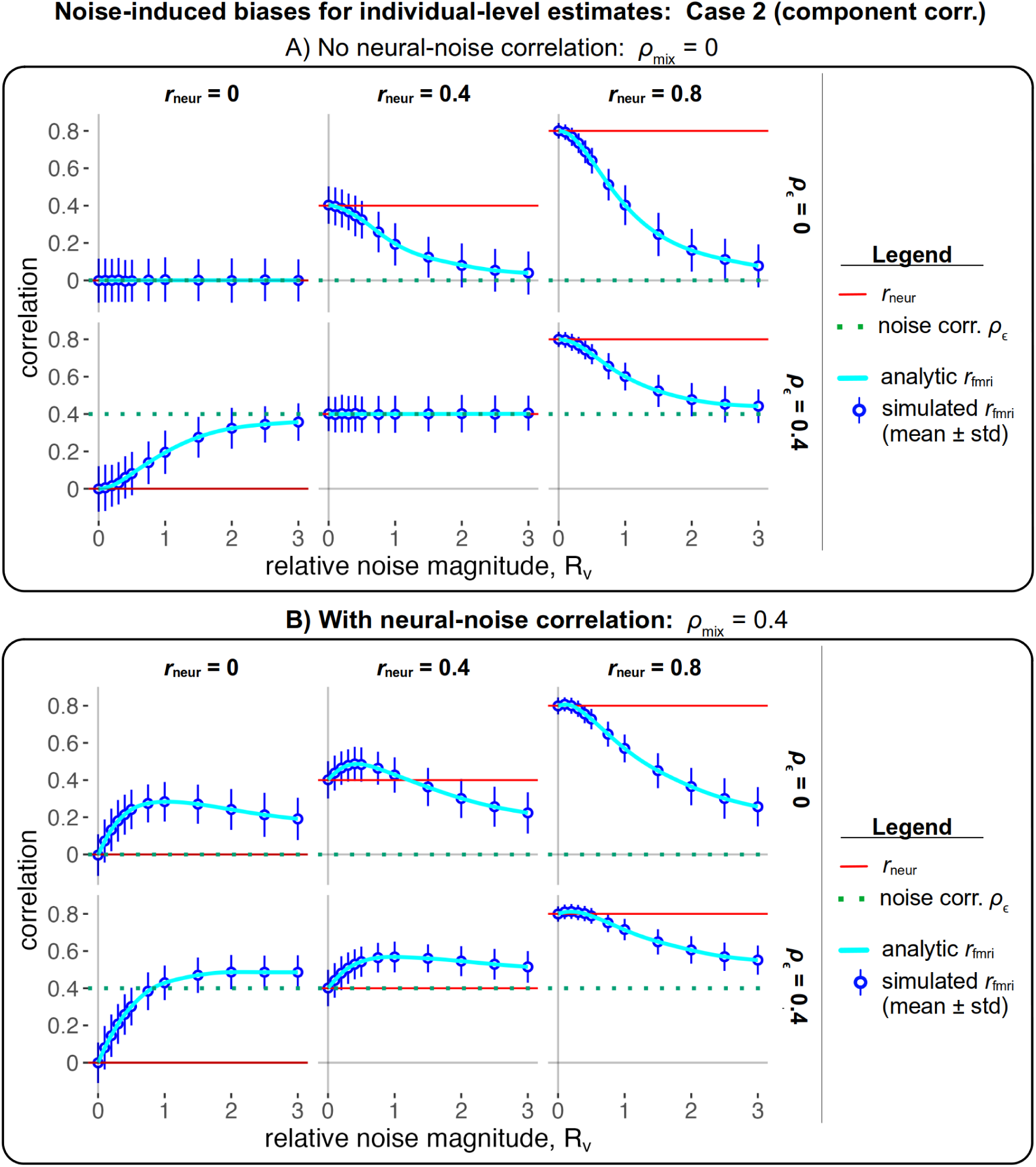
Simulation results illustrating bias in *r*_fMRI_ under neural–noise mixing and shared noise.

Together, these scenarios demonstrate that even moderate and plausible noise conditions can substantially distort correlations. In realistic scenarios with moderate cross- and within-region noise, bias patterns become highly complex: underestimation is common, but overestimation or inflation readily occurs, consistent with empirical observations (Korponay et al., 2024). These difficulties extend to the group level, where resting-state analyses are typically applied, such as in studies of brain development, aging, and disorders including Alzheimer’s disease and schizophrenia (Yamada et al., 2017; Ibrahim et al., 2021). Furthermore, factors that alter the shape of the hemodynamic response can lead to alterations in resting-state fMRI correlations even when the underlying neural correlations remain perfectly identical (Liu, 2013).

### 2.2 Noise-induced bias in correlation estimates: group-level comparisons

Distortion at the individual level rapidly propagates to group-level analyses. Differences in noise structure between groups can influence results in significantly varying ways (Fig. 7): fabricating spurious group differences, suppressing true differences, or even completely reversing the direction of genuine differences. Specifically, variations in noise magnitude, noise synchrony, or neural–noise coupling can generate false group differences (Fig. 7A), whereas high noise levels may obscure (Fig. 7B) or invert genuine differences (Fig. 7C).

**Figure 7:**
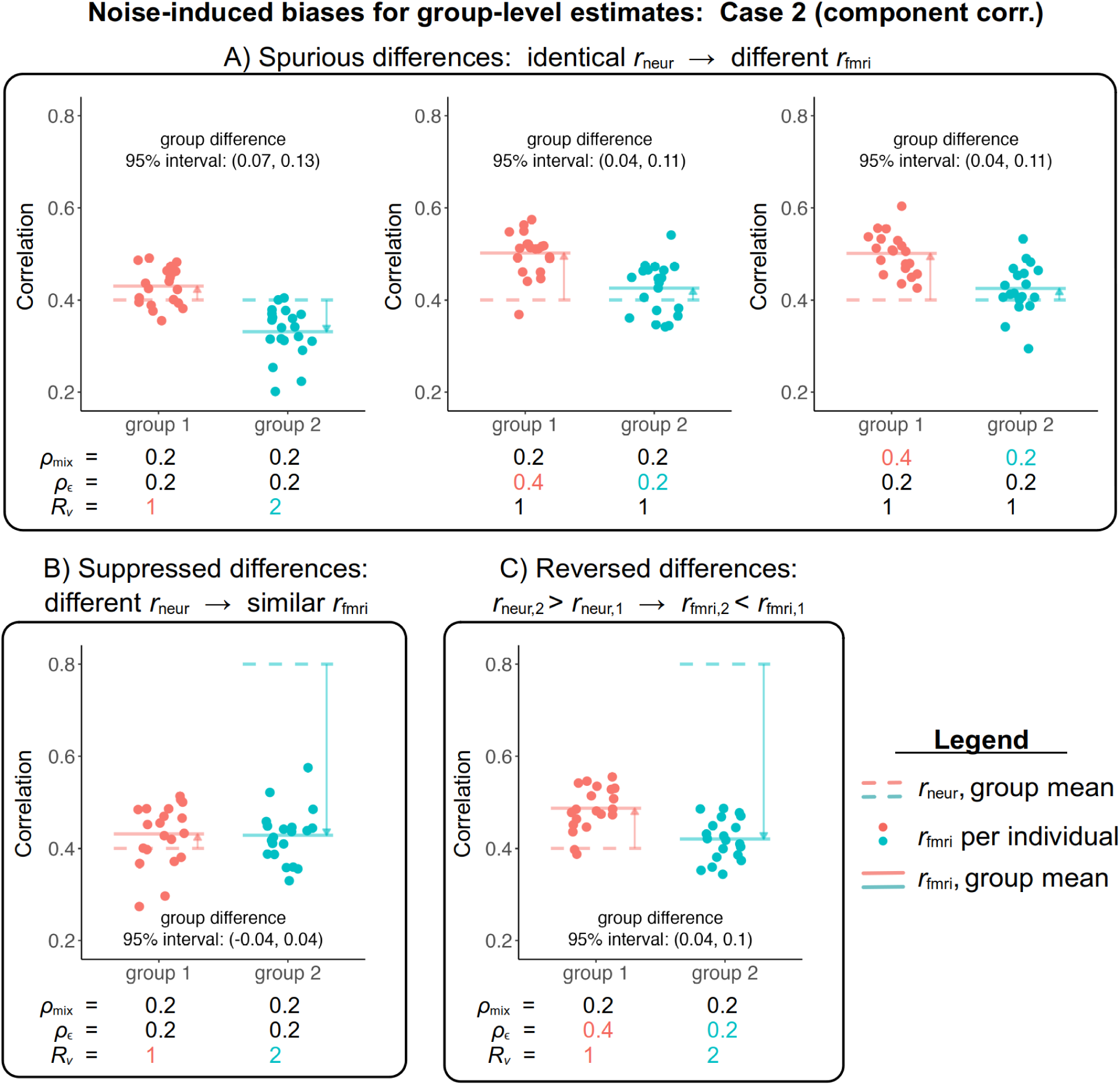
Examples of spurious, suppressed, and reversed group differences arising from noise.

These effects arise naturally from differences in motion, physiology, or vascular properties across populations. While investigators routinely attempt to control noise sources, complete mitigation is rarely feasible. For example, patient and control populations often possess inherently different exogenous noise traits, such as altered breathing rates or head motion. Vascular differences between aging or clinical groups are even more difficult to measure and account for. These distortions are equally relevant to associations with behavioral traits (Taylor et al., 2021) and naturalistic fMRI data, where inter-individual correlations are subject to similar structural biases.

Importantly, these distortions arise at the level of the observed correlations themselves and are not limited to discrepancies between neural and BOLD signals. Although we express distortion relative to a latent neural correlation for conceptual clarity, the same attenuation, inflation, and group-level bias arise whenever the observed correlation is interpreted as a direct estimate of any underlying signal component.

### 2.3 Persistent challenges: the inverse problem of noise

Noise in resting-state fMRI arises from a complex combination of neurovascular coupling, physiological processes, mental states, and measurement-related factors. Importantly, in this context “noise” does not imply purely exogenous or random disturbance, but rather encompasses all variability not explicitly modeled at the neural level, including structured and biologically mediated transformations. While preprocessing techniques such as bandpass filtering, motion censoring, and physiological signal regression can reduce some sources of exogenous noise (Caballero-Gaudes and Reynolds, 2017), complete removal is fundamentally limited. Because the underlying neural activity and the various sources of measurement and physiological variability are not directly observable, denoising methods necessarily rely on measured nuisance variables or proxy estimates (e.g., head motion, respiratory and cardiac recordings, or global signal fluctuations), which are themselves imperfect and subject to measurement error. Simple procedures such as motion censoring can be challenging and controversial (Mejia et al., 2026). Moreover, some nuisance measures partly overlap with genuine neural or neurovascular processes, making complete separation impossible in principle. Global signal removal illustrates this trade-off: although it can reduce widespread physiological and motion-related influences, it may also remove neurally relevant variance and introduce additional distortions (Murphy and Fox, 2017). Consequently, even sophisticated preprocessing cannot uniquely recover the underlying neural correlations from the observed fMRI signal, and the noise scale often remains comparable to or exceeds the neural-related fMRI signal.

Challenges due to the complexity of neurovascular processes are even more problematic, as the spatial variability of neurovascular coupling remains largely unaddressed (Rangaprakash et al., 2023). Thus, current methods cannot fully decouple their influence on the fMRI signal. For instance, the relationship between neural activity and the BOLD responses is demonstrably nonlinear (Zhang et al., 2019), and empirical estimates suggest that neural events explain only a modest fraction of resting-state BOLD variance in some settings (Lu et al., 2019). This establishes a strict practical lower bound on achievable noise reduction.

Physiological and group-related factors exacerbate these challenges. Age-related changes in neurovascular coupling and autonomic function (e.g., heart rate variability) can mimic or obscure developmental effects (Tsvetanov et al., 2020). Motion artifacts, particularly prevalent in populations such as ADHD or Parkinson’s patients, and respiration-induced noise further confound interpretation (Tu and Zhang, 2022). Pharmacological manipulations, including psychedelics, alter vascular dynamics and complicate fMRI signal interpretation (Cardone et al., 2024). These latent sources underscore the need for a more explicit mapping from fMRI correlations to underlying neural activity—one that accounts for spatial and temporal heterogeneity. Without such a framework, correlation-based analyses risk producing spurious or reversed interpretations, limiting their utility for mechanistic biomarker discovery (Taylor et al., 2021; Tu et al., 2024).

An often underappreciated source of distortion arises from mental states such as participant drowsiness and microsleeps. Even during relatively short scans (5–10 minutes), such events can occur multiple times and substantially distort correlation estimates, particularly in default mode and sensory networks (Tagliazucchi and Laufs, 2014). One proposed strategy to avoid this is deriving resting-state-like signals from task-based fMRI by regressing out modeled task responses, baseline, and motion effects. However, this procedure fails to fully capture task-related influences, including mis-specification of the hemodynamic response function, unmodeled cognitive demands, tonic baseline shifts associated with resting-state dynamics (as opposed to phasic task-related events) (Grimm et al., 2024). Consequently, residualization is unlikely to fully isolate true resting-state activity.

These simulations are not intended to reproduce empirically calibrated parameter regimes. Rather, they illustrate a structural property of the measurement process: when neural signals are observed through a noisy and heterogeneous transformation, the mapping from neural activity to observed correlations is not uniquely determined (i.e., multiple underlying configurations can yield identical observed correlations). Because neural-level correlations and noise structure are not directly observable in human fMRI, there is no accessible ground truth (e.g., neural correlations, noise correlations across regions, or their coupling) against which such parameters can be calibrated. Consequently, it is not possible to determine whether a given correlation estimate is attenuated, inflated, or unbiased. This ambiguity reflects the measurement distortion described throughout this section: a fundamental inverse problem at the level of measurement and cannot be resolved from correlation data alone. A related but distinct form of causal non-identifiability arises at the level of causal inference, where even perfectly measured correlations do not uniquely determine underlying interactions (Section 3).

These results imply that, in resting-state fMRI, correlation estimates do not, by themselves, identify causal relationships even at the level of the observed BOLD responses, as the contributions of neural activity and non-neural influences cannot be separated from the data alone. This ambiguity is further compounded by modern large-sample statistical inference. As sample sizes increase to thousands of individuals (Marek et al., 2022), statistical power to detect measurement bias also increases. Trivial but systematic differences in noise variability or neurovascular coupling across groups will generate overwhelming statistical evidence, producing the illusion of a meaningful difference where none exists at the neural level.

In summary, measurement distortion limits how accurately observed correlations represent the underlying processes they are intended to characterize. With multiple underlying neural and physiological configurations producing the same observed correlation structure, there is no unique translation from observed BOLD correlations back to the underlying neural interactions. This does not invalidate correlation-based analysis, but it constrains how directly such correlations can be interpreted as evidence of neural interactions.

## 3 Limits of correlation for causal interpretation

The previous section addressed limitations arising from how correlations are measured. Here we consider a distinct structural limitation, causal non-identifiability: the inability to uniquely infer causal structure from correlation alone. This non-identifiability places fundamental limits on causal interpretation, which is particularly consequential in resting-state fMRI analyses where unanchored correlation matrices serve as the primary mathematical foundation for brain networks.

The ambiguity described in Section 2 arises from the measurement process, where noise and neurovascular variability distort the mapping between neural activity and observed correlations. By contrast, the limitation considered here persists even under idealized conditions with perfect measurement. In this setting, correlation structure itself does not uniquely determine causal relationships. This reflects a distinct form of non-identifiability that is inherent to correlation-based inference and cannot be resolved by improving measurement accuracy without additional assumptions or external constraints.

### 3.1 Non-identifiability in noiseless networks

In practice, resting-state analyses involve networks with many regions, where undirected correlations are used to infer relationships among inherently directed neural processes. It is often implicitly assumed that large positive correlations reflect strong direct interactions, and that sign and magnitude carry straightforward mechanistic meaning. However, even under idealized conditions with perfect measurement and no noise, such interpretations are not uniquely supported by correlation structure (Papo and Buldú, 2025; Zalesky et al., 2012; Rubinov, 2023). To make this limitation concrete, we consider minimal network examples under idealized conditions with linear relationships and perfectly estimated neural-level correlations. If ambiguity persists even under these highly favorable assumptions—linear relationships, acyclicity, and the absence of hidden confounders—it reflects a fundamental mathematical limitation rather than a consequence of measurement error or model complexity. In a two-region system (Fig. 8A), the observed correlation constrains the magnitude of association but does not determine directionality: two equally plausible causal structures exist with opposite directions of influence. In this special case, the magnitude and sign of the association are preserved, but direction remains ambiguous.

**Figure 8:**
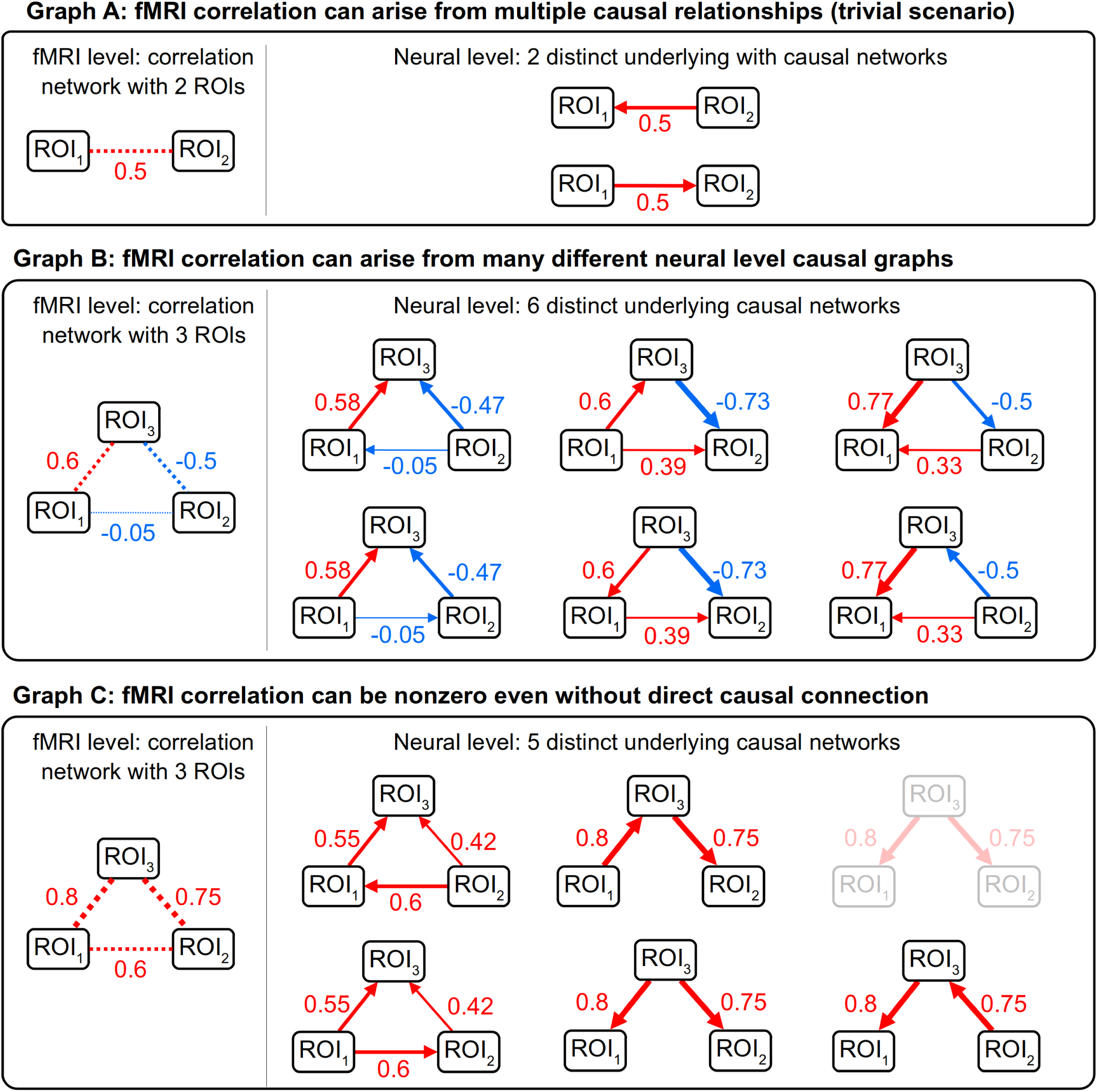
A single observed correlation pattern can arise from multiple underlying causal structures. Dotted lines indicate correlations, solid lines denote directed influences, red/blue indicate positive/negative values, and line thickness reflects magnitude. (A) In a two-region system, correlation constrains the magnitude of association but does not determine directionality. (B) In a three-region system, the same correlation pattern can be generated by multiple distinct causal graphs. (C) Correlation does not imply direct causal link: associations may arise through mediation or confounding (similar examples were heuristically illustrated in Reid et al. (2019)); one of the two indistinguishable causal graphs due to degeneracy is shown faded.

This correspondence breaks down with the introduction of a third region. In general, a set of pairwise correlations among three regions can be generated by multiple distinct acyclic causal graphs (Appendix B). A single correlation pattern may correspond to several causal structures with substantially different strengths of causal influence and even different signs (Fig. 8B). For example, an observed correlation of *r*_fMRI_ = 0.6 between two regions may correspond to true underlying causal strengths of 0.57, 0.6, or 0.77.

More strikingly, a near-zero correlation between two regions can still arise from a substantial direct causal effect, depending on the broader network structure. Even if this correlation were exactly zero, the set of compatible causal structures would persist with only minor adjustments to parameter values. In some cases, strong correlations may arise even in the total absence of any direct causal link between regions (Fig. 8C), reflecting indirect pathways or shared inputs.

This example demonstrates that under ideal conditions, correlation does not uniquely identify either the structure or strength of underlying interactions. More broadly, causal non-identifiability extends beyond uncertainty in causal direction or ordering. Even with a perfectly measured correlation matrix, fundamentally different causal mechanisms—including direct effects, mediation, common-cause structures, and other causal configurations—may generate identical statistical associations. This non-uniqueness is precisely the causal non-identifiability discussed in this section, reflecting a phenomenon known as a *Markov equivalence class*: multiple causal graphs can be observationally equivalent, meaning they imply the exact same set of statistical dependencies despite differing underlying mechanisms. In such cases, the observed data alone are insufficient to distinguish among these competing causal explanations.

These examples clarify the limitations of the three heuristic assumptions introduced in Section 2: zero correlation does not guarantee an absence of interaction, correlation sign does not reliably identify excitatory or inhibitory influence, and correlation magnitude does not cleanly quantify causal strength. These limitations arise from non-identifiability and are independent of measurement noise. This distinction is critical: unlike the distortions discussed in Section 2, these ambiguities persist even under perfect measurement and *cannot* be resolved by improving data quality, refining preprocessing pipelines, or increasing sample size. They represent a hard mathematical boundary on what can be inferred from a correlation matrix. These structural limitations become particularly consequential when correlation matrices are transformed into network representations.

### 3.2 Implications for higher-order representations of correlation structure

Graph-theoretic analyses (Sporns, 2018), gradient-based embeddings (Margulies et al., 2016), and other geometric or topological representations (Pang et al., 2023) summarize correlation structure through deterministic transformations of the underlying associations. As a result, they inherit the same ambiguities and non-identifiability present in the correlations themselves: multiple distinct causal structures can yield identical higher-order representations and derived metrics. Similar considerations apply to latent-state and dynamic network approaches, including Hidden Markov Models, where inferred network states and transitions remain deterministic functions of the same underlying correlation structure and therefore inherit the same measurement distortion and inferential ambiguities.

Accordingly, mechanistic interpretation of derived quantities—including node importance (e.g., degree, centrality, hubness), local structure (e.g., clustering, rich club organization), global organization (e.g., efficiency, density), gradients, latent states, or other geometric and topological representations—as direct evidence of neural mechanisms is structurally underdetermined. These methods may provide useful descriptive summaries, predictive features, or low-dimensional representations of correlation structure, but they cannot resolve the measurement distortion or causal non-identifiability inherent in the underlying correlations. These observations do not diminish the descriptive, predictive, or computational utility of such methods; they do not, by themselves, justify mechanistic interpretation. Rather, they clarify the level of inference that can be supported by the resulting representations.

This ambiguity is exacerbated by common network construction practices. For instance, thresholding weak correlations is a standard step in graph-theoretic analysis. Consider the near-zero correlation (−0.05) in Fig. 8B. At the fMRI level, this weak correlation would typically be removed, interpreted as “no functional connectivity.” However, due to non-identifiability, the same pattern of correlations may still be consistent with a meaningful underlying causal interaction between those regions. Thus, thresholding can remove correlations that correspond to causally relevant relationships, introducing divergence between the estimated correlation network and the underlying causal structure. Conversely, strong correlations that survive thresholding may arise from indirect pathways or shared inputs rather than direct interactions. As a result, thresholding does not resolve ambiguity but instead reshapes it, embedding additional assumptions about which relationships are considered relevant. This illustrates that sparsification procedures operate on statistical association rather than causal structure.

Similarly, other procedures in common practice may preferentially retain biased correlations and discard meaningful ones. For instance, negative correlations are frequently filtered out, set to zero, or transformed via absolute-value operations. These practices largely reflect historical methodological constraints—as many widely used graph metrics were originally defined for non-negative adjacency matrices—rather than principled evidence that negative correlations are uninformative. Such conventions discard data rather than resolving its ambiguity, thereby embedding unverified implicit assumptions about relevance and information flow directly into the network architecture. In addition, sparsity-inducing and feature-selection procedures, including proportional thresholding, LASSO-based selection, and related regularization methods, operate directly on estimated correlations or derived network features. While such approaches may improve statistical stability, interpretability, or predictive performance, they cannot resolve ambiguities inherent in the underlying correlation structure. Consequently, these procedures improve statistical representation rather than mechanistic identifiability; selected features the inferential limitations of the underlying correlation structure, and they cannot exceed the inferential content of its inputs.

A broader concern is the tendency for statistical associations to become reified as mechanistic entities (“connections,” “edges,” “hubs,” “gradients,” “hierarchies”) through terminology and visualization. Correlations are commonly referred to as “functional connectivity” and visualized as network links, chords (e.g., Circos plots), or other spatial representations that resemble physical connections. Such visualizations provide effective descriptive summaries of association structure, but, together with graph-theoretic terminology such as “edges,” “hubs,” “communities,” and “hierarchies,” they may also encourage the perception that statistical associations correspond directly to neural communication pathways or organizational principles. Because these representations remain deterministic transformations of the same underlying correlations, they do not resolve the measurement distortion or causal non-identifiability discussed above. Consequently, these representations should be interpreted as increasingly sophisticated summaries of correlation structure, not as progressively stronger evidence of underlying neural mechanisms.

Moreover, graph-based analyses typically do not propagate uncertainty from the underlying correlations. As a result, quantities such as centrality, modularity, and efficiency may appear highly stable and interpretable, while actually reflecting ambiguous and potentially non-unique underlying structures. This lack of standard errors can lead to the overconfidence and overinterpretation of network patterns as reflecting true neural communication or organization.

The limitations demonstrated above were shown for the simplest possible scenario: three regions with perfect, noise-free neural-level correlations. These results indicate that interpretive ambiguity is not merely a product of noisy fMRI data; it is inherent to the correlational framework itself. In realistic whole-brain networks involving tens or hundreds of regions, the number of compatible causal configurations increases combinatorially with network size. That is, the difficulty of recovering a unique mapping from the BOLD correlation level to the causal strength at the neural level dramatically and rapidly amplifies.

In summary, non-identifiability is a structural limitation of correlation-based analysis: even if measurement distortion were fully resolved, the mapping ambiguities from causal structure to correlation would remain. In practice, this limitation is inextricably compounded by the measurement distortion, nonlinear dynamics, and latent confounding discussed in Section 2. Together, these factors imply that correlation-based analyses provide descriptive summaries of association structure, but do not, by themselves, support mechanistic inference about neural interactions.

## 4 Discussion

Resting-state fMRI has reshaped systems neuroscience by revealing large-scale patterns of coordinated brain activity that are reproducible across individuals, populations, and acquisition settings. These patterns have informed models of functional organization, development, and disease, and have supported clinically relevant applications. At the same time, correlation-based analyses provide observational summaries of BOLD time series rather than direct measurements of neural interaction. Like all measurement systems, they have limits that must be considered when interpreting results.

The present work does not challenge the descriptive or predictive value of resting-state fMRI. Rather, it clarifies the inferential boundaries governing its interpretation. In other words, correlation is informative, but not mechanistically self-interpreting. In particular, we distinguish three levels of inference (Table 2). First, *descriptive association*: correlations summarize statistical dependence in measured signals and are valid at that level, though subject to bias from measurement distortion. Second, *predictive or biomarker utility*: reproducible correlation patterns can be used effectively for classification, prediction, and intervention targeting, even when influenced by non-neural factors; however, predictive performance may be sensitive to measurement bias and does not imply identification of underlying neural mechanisms. Third, *mechanistic or causal interpretation*: claims about neural interaction, directionality, or information flow are not identifiable from correlation structure alone and require additional assumptions or experimental constraints.

**Table 2:** Levels of interpretation for correlation-based resting-state fMRI analyses.

| Level | Interpretation | What is supported / limitations |
| --- | --- | --- |
| Descriptive association | Correlation summarizes statistical association between fMRI signals | Supported as a property of measured signals; may be biased by measurement distortion, including spurious, suppressed, or sign-reversed group differences |
| Predictive / biomarker utility | Correlations used for classification, prediction, or intervention targeting | Supported even with non-neural contributions. Predictive performance may remain strong despite measurement distortion, but feature importance does not imply mechanistic importance |
| Mechanistic / causal interaction | Correlations interpreted as neural interaction or information flow | Not identifiable from correlation alone; requires additional assumptions or data |

### 4.1 Consequences of measurement distortion

Measurement distortion arises because the fMRI signal is an indirect and heterogeneous transformation of neural activity. As shown in Section 2, plausible noise structures can attenuate, inflate, or otherwise bias estimated correlations, including at the group level. These effects arise from the measurement process itself and are not eliminated by increasing sample size. As a result, precisely estimated correlations may still be systematically biased. Correlations are computed on signals that inextricably mix neural and non-neural influences. Thus, the issue is not only how correlation is interpreted, but also that the quantity being analyzed is itself a distorted proxy of neural activity. Consequently, there is no unique translation from the measured correlation structure to the underlying neural processes.

Because neural-level correlations and noise structure are not directly observable, these ambiguities cannot be empirically resolved in human fMRI data alone. Thus, the issue is not only how correlation is interpreted, but also that the quantity being analyzed is itself a distorted proxy of neural activity. Moreover, these limitations arise at the level of the observed BOLD responses itself. Consequently, they apply regardless of whether inference targets neural activity or hemodynamic dynamics, as the observed correlations reflect mixtures of multiple underlying processes rather than direct estimates of any single component.

These considerations can also be framed in terms of classical notions of validity. The indirect and heterogeneous mapping from neural activity to BOLD responses limits construct validity, as correlation-based measures do not uniquely correspond to specific neural processes. Measurement distortion and latent confounding further constrain internal validity, as correlation structure alone does not permit causal interpretation. At the same time, resting-state fMRI often demonstrates strong reproducibility across datasets and populations, reflecting a form of external validity. These dimensions are not equivalent: reproducibility of correlation patterns does not imply accurate measurement of neural processes or valid causal inference.

A key implication concerns reproducibility. Correlation patterns are often highly reproducible across studies, scanners, and populations. However, reproducibility reflects the statistical stability of the measured signal, not necessarily its accuracy with respect to underlying neural relationships. Systematic biases—arising from unmodeled neurovascular coupling, physiological confounds, or preprocessing choices—can themselves be highly reproducible. As sample sizes grow, these systematic biases will yield statistically robust, reproducible patterns that are biologically spurious. Thus, reproducibility should not be interpreted as evidence of biological validity without additional support.

These considerations are particularly relevant for large-scale brain-wide association studies (BWAS), where increasing sample size improves statistical sensitivity for detecting stable association patterns but does not resolve measurement distortion or causal non-identifiability. As a result, highly reproducible associations may support discovery and prediction while still reflecting mixtures of neural and non-neural influences. Recent large-scale BWAS have demonstrated robust associations between socioeconomic and neuroimaging variables across populations. Such findings exemplify the distinction between predictive utility and mechanistic interpretation: reproducible associations may be informative and clinically useful without uniquely identifying the underlying neural mechanisms. More generally, statistical evidence, predictive utility, effect size, and causal importance are distinct concepts. Large sample sizes can yield overwhelming statistical evidence even for modest associations, and highly reproducible or predictive findings need not correspond to large causal effects. Consequently, rankings based on statistical evidence, reproducibility, or predictive performance should not be interpreted as rankings of causal importance or mechanistic influence.

### 4.2 Consequences of causal non-identifiability

Even in the total absence of measurement distortion, correlation structure does not uniquely determine causal architecture. As shown in Section 3, the mapping from causal structure to correlation is many-to-one. Multiple distinct causal configurations can generate identical correlation matrices, meaning that the inverse problem of recovering neural interactions from correlations is fundamentally underdetermined. From this perspective, the term “functional connectivity,” while historically entrenched, may inadvertently encourage interpretations that exceed what correlation structure alone can support.

This limitation is structural rather than statistical: it cannot be resolved by larger sample sizes, cleaner data, or improved estimation precision. Correlation is symmetric and does not distinguish direct from indirect effects, nor does it uniquely encode the sign or strength of causal interactions. As a result, mechanistic interpretations of correlation-based analyses strictly require additional assumptions, external perturbations, or complementary data. Together with measurement distortion, this structural non-identifiability tightly constrains the extent to which correlation matrices can support mechanistic conclusions.

### 4.3 Implications for analysis and interpretation

Measurement distortion and causal non-identifiability imply that there is no unique translation from observed resting-state correlation structure to the underlying neural interactions it is often intended to represent. These limitations have direct implications for common analytical practices. Graph-theoretic analyses, which transform correlation matrices into macroscale network representations, inherit both measurement distortion and non-identifiability. Because network metrics such as centrality, modularity, and efficiency are deterministic functions of the underlying matrix, they summarize statistical associations but cannot uniquely specify underlying neural circuitry. Similarly, gradient-based representations provide elegant, low-dimensional summaries of correlation structure, but remain subject to the same fundamental ambiguities. Deterministic transformations, dimensionality reductions, latent-state decompositions, and feature-selection procedures cannot create inferential information absent from the original correlation structure. This observation concerns interpretation rather than utility.

These issues are particularly consequential in resting-state fMRI because correlation structure itself is the primary object of inference. In task-based paradigms, experimental manipulation introduces externally controlled variation that provides an inferential anchor: regression models relate BOLD responses to known inputs, allowing task-related signals to be partially separated from exogenous noise. This separation is further supported by experimental design features such as randomization of trial conditions and temporal jittering, which help reduce systematic coupling between task effects and physiological or measurement-related variability. By contrast, resting-state analyses lack such design-based constraints, and these influences are absorbed directly into the estimated correlations.

Given these constraints, stronger interpretability may be achieved through complementary approaches that provide additional anchors or improve modeling of neurovascular coupling. Multimodal integration (e.g., vascular imaging), perturbation-based methods (e.g., TMS–fMRI), high-resolution temporal measurements (e.g., EEG/MEG), structural connectivity references (e.g., diffusion tractography), and models that explicitly incorporate physiological structure can provide necessary constraints on causal inference.

For example, early findings of reduced resting-state correlation due to caffeine were unable to determine whether the effects were driven primarily by neural or vascular mechanisms (Wong et al., 2012). The use of simultaneous EEG measurements helped establish that caffeine-related changes in arousal led to a decrease in the global signal, accounting for both the reduction in estimated correlations and the observed changes in network organization (Wong et al., 2013). Subsequent multimodal studies have further clarified how arousal modulates global brain activity and correlation structure (Liu and Falahpour, 2020).

It is generally believed that brain structure and function are fundamentally linked. However, no single modality provides a complete characterization of neural interactions. For example, structural connectivity derived from diffusion tractography and correlation patterns derived from resting-state fMRI exhibit only partial correspondence (Straathof et al., 2018; Liu et al., 2023; Popp et al., 2024), reflecting the fact that anatomical pathways and statistical cofluctuations capture different aspects of brain organization. This incomplete correspondence is consistent with the broader challenges discussed in this paper, namely that measurement limitations and inferential non-identifiability can complicate the mapping from observed signals to underlying neural interactions. Because structural and functional patterns characterize different aspects of brain organization through distinct measurement processes, correspondence between them should not be interpreted as a direct test of mechanistic validity. Instead, convergence across complementary modalities provides stronger support for mechanistic interpretations than any individual modality alone.

Predictive or biomarker utility does not imply that the underlying features accurately reflect neural interactions. Measurement distortion can introduce systematic biases into the features used for prediction, such that predictive performance may partly rely on stable non-neural factors (e.g., physiological or vascular signals). Because the observed correlation structure is itself a systematically distorted representation of the underlying neural interactions, predictive models necessarily learn from these distorted features. Consequently, the estimated predictive importance of correlation-based features may differ substantially from the importance of the underlying neural processes they are intended to represent. Predictive models therefore rank the informativeness of the measured features, not necessarily the importance of the underlying neural mechanisms. In addition, causal non-identifiability implies that predictive features need not correspond to unique underlying neural mechanisms. An additional distinction concerns the interpretation of predictive models. Predictive utility concerns the performance of the model, whereas mechanistic interpretation concerns the meaning of the predictors. These are distinct inferential questions. High predictive performance establishes that a variable contributes useful information for prediction, but it does not, by itself, establish why that variable is predictive. Likewise, changes in predictor importance following adjustment for additional variables should not be interpreted directly as evidence of causal mechanisms without additional assumptions regarding confounding, mediation, or the underlying causal structure. In other words, predictive importance is not equivalent to mechanistic importance. The legitimacy of prediction is not in question; rather, interpreting what makes a prediction work is a separate scientific problem, subject to the same measurement distortion and causal non-identifiability discussed throughout this paper.

### 4.4 Looking forward

Scientific investigation ultimately seeks explanations of underlying mechanisms and causes. Consequently, statistical associations naturally invite causal interpretation, particularly when they are strong, reproducible, or predictive. This tendency is reflected not only in causal language (e.g., “matter,” “drive,” “shape”), but also in terminology and visual representations that reify statistical associations as mechanistic entities—for example, networks, hubs, communities, gradients, or topological organization. However, these descriptions may imply a level of mechanistic understanding that exceeds what the underlying data alone can support. The transition from association to mechanism generally requires assumptions and evidence that extend beyond the association structure itself, regardless of whether that structure is expressed as correlations, networks, gradients, latent states, or other higher-order representations.

More generally, large-scale association studies often generate debates that appear to concern substantive scientific mechanisms, when they partly reflect differences in the level of inference supported by the data. Likewise, the notion that a brain region, network, or explanatory variable is important is itself ambiguous. Importance may refer to strong statistical evidence, large effect size, predictive utility, graph-theoretic prominence, or causal influence. These concepts address fundamentally different scientific questions and should not be interpreted interchangeably. Distinguishing among these concepts—and, more generally, among descriptive association, predictive utility, and mechanistic interpretation—helps clarify what correlation-based analyses can reveal directly and what requires additional assumptions or complementary evidence.

The limitations arising from measurement distortion and causal non-identifiability can also be viewed through the lens of classical validity concepts (Shadish et al., 2002). Measurement distortion and the indirect nature of the BOLD responses limit construct validity, as correlation-based measures do not uniquely correspond to specific neural processes. Causal non-identifiability constrains internal validity, as correlation structure alone cannot establish directional or mechanistic relationships. At the same time, resting-state fMRI often demonstrates high reproducibility across datasets and populations, reflecting a form of external validity. Importantly, these dimensions need not align: correlation patterns may be highly reproducible while remaining limited in construct validity and non-identifiable for causal inference.

A central implication of this work is the need to more clearly define the target of inference in resting-state fMRI. Correlation-based analyses provide well-defined statistical summaries, but the underlying neural quantities they are intended to represent are too often left implicit. Explicitly defining these targets helps align analytic methods with scientific questions and clarifies what additional assumptions are required for mechanistic interpretations.

None of these limitations diminish the empirical contributions of resting-state fMRI. Correlation patterns capture meaningful aspects of brain organization and continue to serve as highly powerful predictive biomarkers. The key point is that the leap from statistical association to biological mechanism requires assumptions that go far beyond the correlation matrix alone. Recognizing this distinction is not a critique of the methodology, but a prerequisite for its responsible and effective scientific use.

## 5 Conclusions

Correlation-based measures in resting-state fMRI provide reproducible and descriptively powerful summaries of large-scale brain organization. Their interpretational limits arise from two distinct, structural constraints: *measurement distortion*, reflecting the indirect and physiologically mediated nature of the fMRI signal, and *causal non-identifiability*, reflecting the many-to-one mapping from neural causal structure to statistical correlation. Together, these constraints strictly delimit what can be inferred about neural processes from correlation matrices alone, regardless of sample size or statistical precision.

These limitations do not diminish the utility of resting-state fMRI, but they do clarify the levels at which its results can be safely interpreted. Correlations are well-suited for descriptive and predictive purposes, such as biomarker development and classification. However, mechanistic claims regarding information flow or circuit organization require additional anchors, such as external perturbations or multimodal constraints. This concerns interpretive scope rather than methodological utility.

Future progress will likely depend on strengthening these inferential anchors through biophysically informed modeling, longitudinal designs, and the integration of complementary modalities. Yet, even significant methodological refinements will not collapse the fundamental distinction between statistical association and causal structure. BOLD correlations encode meaningful information about brain organization, but the mapping to neural mechanisms remains an underdetermined inverse problem. Recognizing these limits is not a retreat from resting-state research, but a necessary prerequisite for more precise and mechanistically grounded neuroscientific interpretation.

## Acknowledgments

PT and GC were supported by the NIMH Intramural Research Program (ZICMH002888) of NIH/HHS, USA. JF and PAB were supported by NIMH IRP (ZIAMH002783) of NIH/HHS, USA. ZC was funded by the FRQS Postdoctoral Fellowship (doi.org/10.69777/336081), Fonds de recherche du Québec – Santé (FRQS) and the MNI Jeanne Timmins Costello Fellowship, Canada.

This research was supported in part by the Intramural Research Program of the National Institutes of Health (NIH). The contributions of the NIH authors were made as part of their official duties as NIH federal employees, are in compliance with agency policy requirements, and are considered Works of the United States Government. However, the findings and conclusions presented in this paper are those of the authors and do not necessarily reflect the views of the NIH or the U.S. Department of Health and Human Services.

## Appendices

### A Measurement distortion: models and analytic results

This appendix formalizes the measurement models underlying Section 2 and derives analytic expressions for bias in correlation estimates under structured noise.

For a sample of size *n* with true correlation *r*, the expected value of the Pearson estimator is approximately:

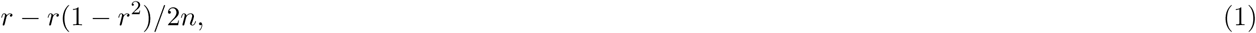

with standard error:

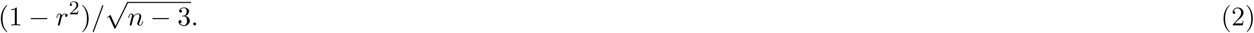

These expressions indicate that sampling bias is negligible for typical fMRI time series lengths, implying that the distortions analyzed in Section 2 arise primarily from measurement structure rather than finite-sample effects.

Parameters and symbols for models considered below are summarized in Table 3.

**Table 3:** Reference table of symbols used in model formulas and simulations.

| Term | Description |
| --- | --- |
| HDR.neural<sub>*i*</sub> | Neural component of the hemodynamic response in region *i* |
| $\sigma_{\text{neur}}$ | Standard deviation of HDR.neural signal |
| $\epsilon_i$ | Noise component of the fMRI signal in region *i* |
| $\sigma_\epsilon$ | Standard deviation of noise |
| fMRI<sub>*i*</sub> | Observed fMRI signal combining neural and noise components |
| $R_v$ | Variability ratio: $\sigma_\epsilon/\sigma_{\text{neur}}$ |
| $r_{\text{neur}}$ | True neural-level correlation |
| $r_{\text{fMRI}}$ | Observed fMRI correlation |
| $\rho_{\text{mix}}$ | Neural–noise mixing correlation |
| $\rho_\epsilon$ | Noise–noise correlation |

#### A.1 Case 1: independent additive noise

Under the assumptions of Fig. 3, we model:

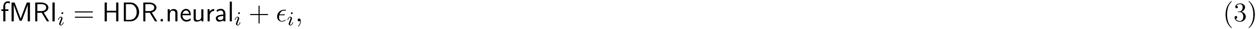

with

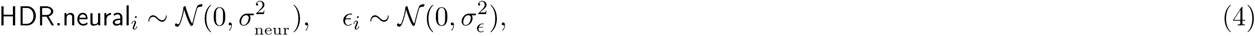

where noise is independent across regions and uncorrelated with neural signals. The resulting correlation is:

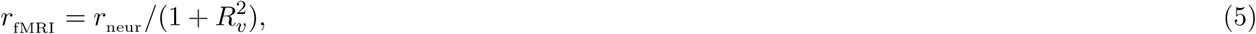

where *R_v_* = *σ_ɛ_/σ*_neur_ quantifies noise relative to the neural signal. Thus, independent additive noise deterministically attenuates correlation toward zero.

#### A.2 Case 2: structured noise and neural–noise coupling

To incorporate more realistic dependencies as shown in Fig. 5, we model the joint distribution:

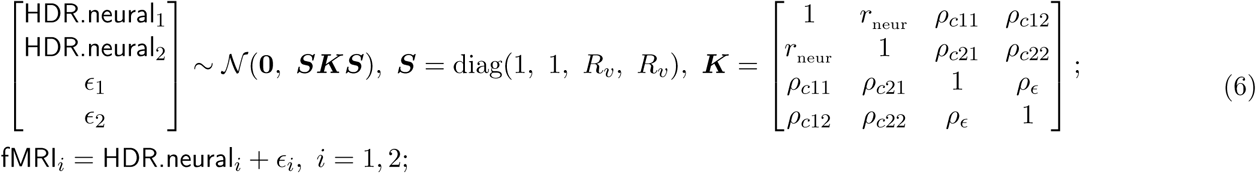

where ***K*** encodes correlations between neural and noise components, and ***S*** scales their variances. The noise term *ɛ_i_* is assumed to contain both exogenous and endogenous sources of measurement distortion. Here, “exogenous” and “endogenous” are used in a causal sense to distinguish influences external to the neural system from variability arising along the measurement pathway. Without loss of generality, we assume zero means for all four variables and a standard deviation of one for the neurally-evoked BOLD responses.

To simplify the complexity of the correlation matrix ***K***, we introduce the assumption of homogeneity among the four correlations between neurally-evoked BOLD responses and noise: *ρ_c_*_11_ = *ρ_c_*_12_ = *ρ_c_*_21_ = *ρ_c_*_22_ = *ρ*_mix_. The correlation *r*_fMRI_ of the fMRI signal between the two regions can be derived as,

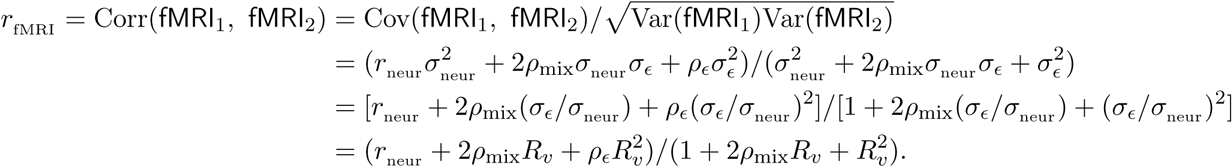

This expression shows that:

1. correlated noise can inflate or induce correlations;
2. neural–noise coupling can alter both magnitude and sign;
3. bias arises as a deterministic consequence of measurement structure.

Thus, measurement distortion reflects not only noise magnitude but also structural dependencies between neural and non-neural processes. These results demonstrate that multiple combinations of neural and noise parameters can yield identical observed correlations. Consequently, the mapping from neural activity to BOLD correlation is not one-to-one, even before considering causal inference. This ambiguity motivates the analysis of structural non-identifiability in Appendix B.

## B Causal non-identifiability among three regions

We demonstrate analytically that a fixed set of three pairwise correlations among three regions does not uniquely determine their underlying causal structure. Importantly, this non-identifiability persists even under highly favorable assumptions, including linear relationships, acyclicity, and the absence of hidden confounders. Multiple distinct causal configurations can generate identical correlation matrices despite these simplifying assumptions. Thus, correlation data alone are insufficient for identifying directionality or causal structure. The assumptions of linearity, acyclicity, and causal sufficiency are adopted only to simplify the analysis. Relaxing these assumptions generally increases, rather than decreases, the space of observationally compatible causal models.

Let *X*, *Y*, and *Z* denote standardized random variables (zero mean, unit variance) representing neural-level signals from three regions, with pairwise correlations *r_xy_*, *r_xz_*, and *r_yz_*, none equal to ±1. We show how one admissible acyclic data-generating process produces these correlations, and note that alternative causal orderings—related by variable permutation—produce the same observable second-order statistics.

The six acyclic orderings on three variables depicted in Fig. 8B share an identical fully connected skeleton and therefore form a Markov equivalence class. Therefore, we illustrate the derivation only for the case in Fig. 9B that corresponds to the first scenario on row 1 of Fig. 8B, while all other cases can be similarly derived through variable rotations. Our goal is to express the causal effects *c_xy_*, *c_xz_*, and *c_zy_* among the three variables in terms of their pairwise correlations *r_xy_*, *r_xz_*, and *r_zy_*.

**Figure 9:**
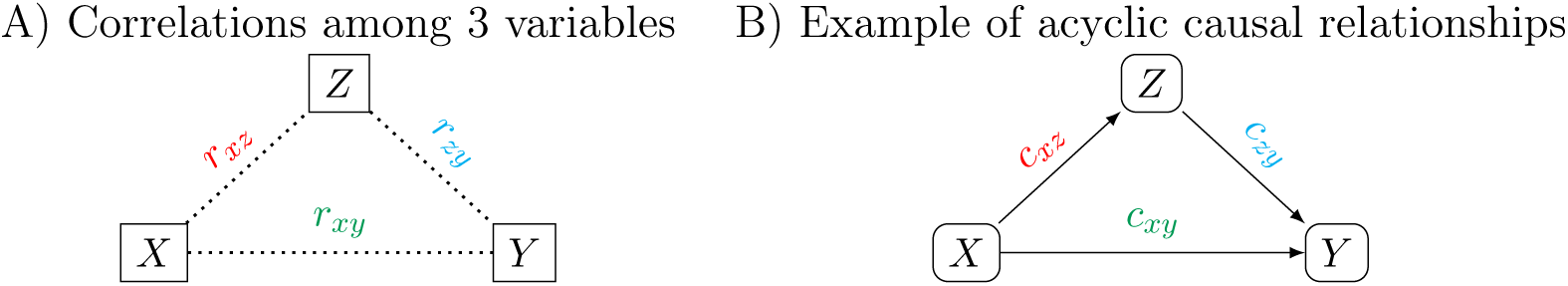
Mapping pairwise correlations to causal relationships. (A) Pairwise correlations among three standardized variables *X*, *Y*, and *Z*: *r_xy_*, *r_xz_*, and *r_zy_*. (B) One example of an acyclic causal configuration consistent with these correlations, with *Z* mediating the effect on *Y* and *X* acting as a confounder for *Z Y* . The causal parameters *c_xy_*, *c_xz_*, and *c_zy_* correspond to the underlying causal effects that can produce the observed correlations.

We construct a model for the linear relationships based on the data-generating process depicted in Fig. 9B,

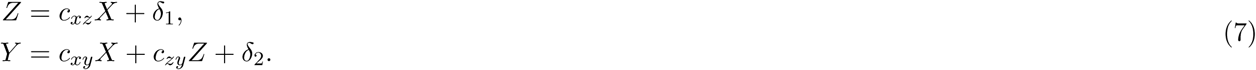

where the zero-mean “noise” *δ*_1_ is independent of *X* while the zero-mean “noise” *δ*_2_ is independent of *X* and *Z*. The parameter *c_xy_* represents the direct causal effect of *X* on *Y*, controlling for the mediator *Z*, while *c_zy_* captures the causal effect of *Z* on *Y*, accounting for the influence of the confounder *X*. With the relationships below for the model (7),

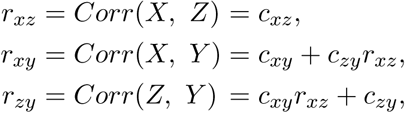

we obtain the following causal effects,

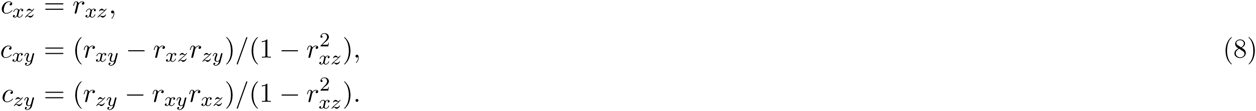

The causal effects illustrated in Fig. 8B-C are calculated using the formula (8). Notably, this indeterminacy

arises even in an idealized linear-Gaussian setting without measurement error or preprocessing artifacts. The ambiguity is therefore structural, not methodological.

### Proposition

Let Σ be a positive-definite 3 × 3 correlation matrix with all pairwise correlations nonzero and strictly between -1 and 1. Then:

i. There exists a linear acyclic structural equation model generating Σ.
ii. Multiple distinct acyclic causal orderings of {*X, Y, Z*} generate the same Σ.

### Implication

Formula (8) shows that, for any admissible correlation matrix, one can construct a consistent acyclic causal parameterization. However, alternative orderings—obtained by permuting the roles of *X*, *Y*, and *Z*—produce identical second-order statistics while assigning different directional interpretations to the effects. When all pairwise correlations are nonzero, the fully connected three-node graph contains no conditional independencies, and the corresponding acyclic orientations form a Markov equivalence class. Consequently, directionality is not identifiable from correlations alone. Additional structure—such as temporal precedence, non-Gaussianity, or experimental intervention—is required to resolve the ambiguity. In other words, the ambiguity is not a consequence of measurement distortion or preprocessing, but arises even in an idealized linear-Gaussian setting.

